# Intersectin1 promotes clathrin-mediated endocytosis by organizing and stabilizing endocytic protein interaction networks

**DOI:** 10.1101/2024.04.22.590579

**Authors:** Meiyan Jin, Yuichiro Iwamoto, Cyna Shirazinejad, David G. Drubin

## Abstract

During clathrin-mediated endocytosis (CME), dozens of proteins are recruited to nascent CME sites on the plasma membrane. Coordination of endocytic protein recruitment in time and space is important for efficient CME. Here, we show that the multivalent scaffold protein intersectin1 (ITSN1) promotes CME by organizing and stabilizing endocytic protein interaction networks. By live-cell imaging of genome-edited cells, we observed that endogenously labeled ITSN1 is recruited to CME sites shortly after they begin to assemble. Knocking down ITSN1 impaired endocytic protein recruitment during the stabilization stage of CME site assembly. Artificially locating ITSN1 to the mitochondria surface was sufficient to assemble puncta consisting of CME initiation proteins, including EPS15, FCHO, adaptor proteins, the AP2 complex and epsin1 (EPN1), and the vesicle scission GTPase dynamin2 (DNM2). ITSN1 can form puncta and recruit DNM2 independently of EPS15/FCHO or EPN1. Our work redefines ITSN1’s primary endocytic role as organizing and stabilizing the CME protein interaction networks rather than a previously suggested role in initiation and provides new insights into the multi-step and multi-zone organization of CME site assembly.

## Introduction

Clathrin-mediated endocytosis (CME) is an essential process in eukaryotes that regulates cell signaling and mediates nutrient uptake, with over 60 proteins involved^1–3^. However, how endocytic protein recruitment is coordinated in time and space is not fully understood.

The CME process consists of initiation, stabilization, maturation, vesicle scission, and uncoating stages (Figure 1A)^3,4^. In the past decade, our understanding of the molecular mechanism of CME site initiation has significantly improved thanks to studies from multiple research groups. Recent studies suggest that endocytic pioneer proteins EPS15^5^ and FCHOs^6–8^ might form liquid-like condensates on the plasma membrane^9^. Previous work suggests that the AP2 complex, the main CME cargo adaptor, is recruited to and activated at nascent CME sites through transient interactions with EPS15 and FCHO^7,8,10,11^. Combining knowledge of the ultrastructural organization of endocytic proteins from super-resolution imaging^12^ and evidence for liquid condensate formation, a two-phase CME protein network assembly model has recently been proposed^13^. This model suggests that subsequent to CME site initiation, proposed to involve the first liquid-liquid phase separation event, a second liquid-liquid phase separation event occurs involving Epsins and CALM, producing a protein network located at the center of the initial condensate, outcompeting EPS15 for binding to the cargo-bound AP2 complex^10,13,14^. However, the molecular mechanism underlying the formation of this second phase and how the two putative phases are connected remains to be determined.

**Figure 1.**
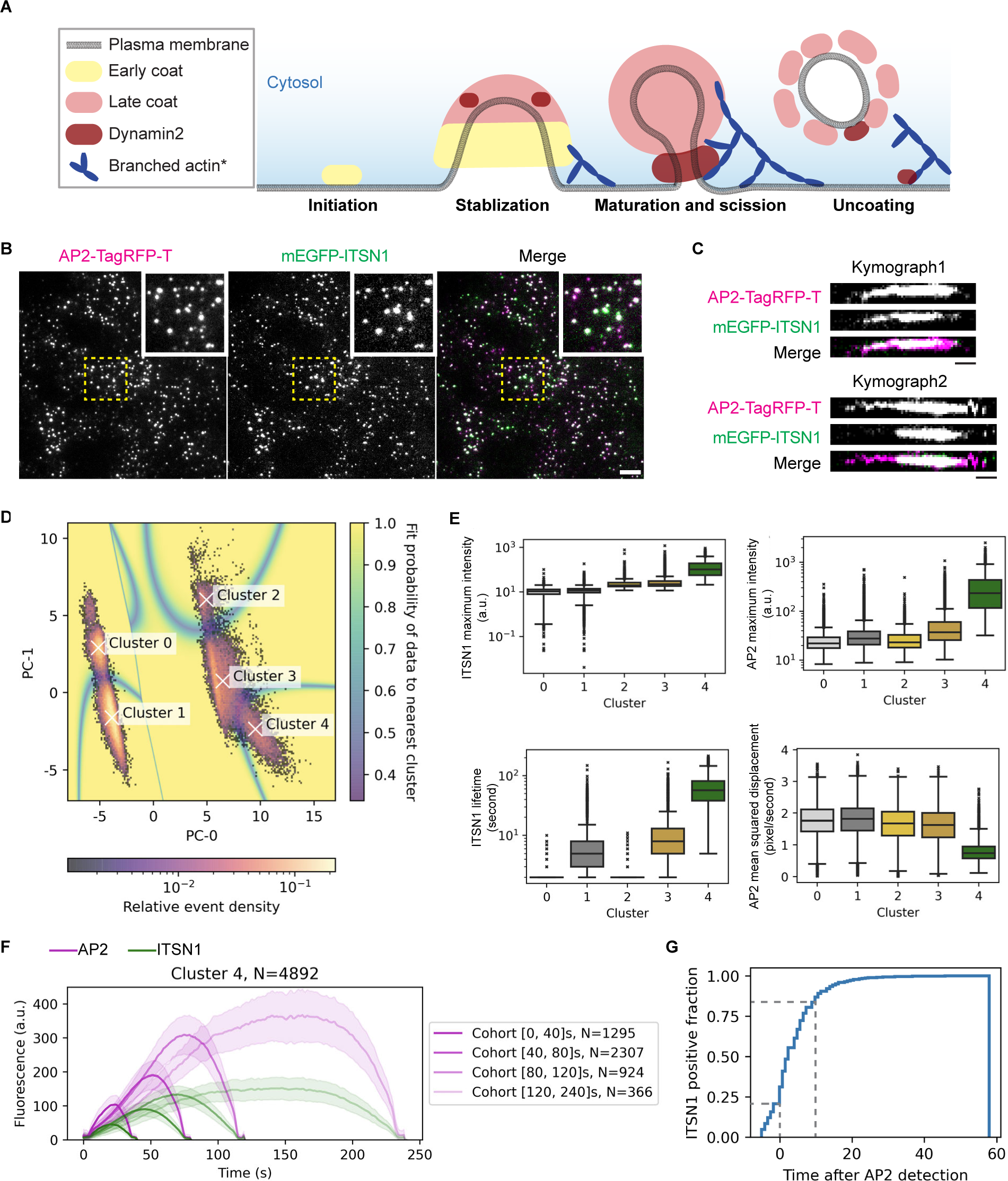
ITSN1 is recruited during the stabilization stage of CME site assembly. (A) Schematic model highlighting different stages of CME referred to in the text. (B) A representative single time frame image from a TIRF movie of AP2M1-tagRFP-T (magenta) and mEGFP-ITSN1 (green) genome-edited human iPSCs. The regions enlarged in the insets are boxed by the dashed lines. Scale bar: 5 µm. (C) Representative kymographs of CME sites in AP2M1-tagRFP-T (magenta) and mEGFP-ITSN1 (green) genome-edited human iPSCs. Scale bar: 10 sec. (D) 2-D histogram of the first two principal components (PCs) of AP2 and ITSN1 dynamic features. Valid tracks detected by cmeAnalysis^24^, appearing and disappearing during movie acquisition were selected for the clustering. Relative event density shows the distribution of the valid tracks in the principal component space, and the shaded underlay (fit probability of data to the nearest cluster) represents the probability of the simulated data points belonging to the nearest cluster. (E) Quantification of ITSN1 and AP2 maximum intensity, AP2 lifetime and mean squared displacement of valid tracks in different clusters. For boxplots, the box extends from the first quartile to the third quartile of the data, with a line at the median. Whiskers extend 1.5x the interquartile range (IQR) from the box. Outliers are shown as “x” marks. Cluster 0: N=9657, Cluster 1: N=24513, Cluster 2: N=2242, Cluster 3: N=13381, Cluster 4: N=4892. (F) Averaged intensity vs time plots for cohorts of Cluster 4 tracks. Tracks are grouped into cohorts by AP2 lifetimes. Numbers of tracks in each cohort are shown next to the plot. Error bar: ¼ standard deviation. (G) Cumulative histogram of the fraction of ITSN1-positive Cluster 4 tracks at a given time after the AP2 signal is first detected. N=4892.

Intersectin1 (ITSN1) is a multivalent scaffold protein known to interact with multiple endocytic proteins^6,15^. ITSN1 has two main isoforms: a short, 140 KDa, universally expressed isoform, ITSN1-S, and a long, 200 KDa, neuron-specific isoform, ITSN1-L, with ITSN1-S being more extensively studied^15–17^. ITSN1’s interactions implicate it as a key protein functioning at the nexus of initiation components (EPS15 and FCHO2) and the centrally-localized proteins (CALM and Epsins). A knockdown screen involving 67 CME proteins suggested that intersectin1 might participate in CME site stabilization^18^. The combined evidence identifies ITSN1 as a candidate to form connections within CME protein networks and bridge putative liquid phases.

In contrast to a proposed role in CME stabilization, ITSN1 has also been proposed to be a CME site initiation complex member because when overexpressed, it arrives at CME sites simultaneously with EPS15 and FCHO, raising questions about whether its true function is in CME site initiation or stabilization^19^. Overexpressed proteins often display altered dynamics and functions^20,21^, however, the timing of endogenously expressed ITSN1’s arrival and its potential role in CME site initiation have not been reported.

In this study, through the combination of genome editing, live cell imaging, high-throughput computational analysis, and a mitochondria relocalization assay, we demonstrate that ITSN1-S plays a crucial role in CME site stabilization by establishing an interaction network that interconnects endocytic proteins that arrive at different stages of CME and localize to different spatial zones within CME sites.

## Results

### ITSN1 is recruited during the stabilization stage of CME site assembly

Previous studies showed that recruitment of the AP2 adaptor complex to CME sites occurs shortly after CME site initiation^6,14^, a process that some studies suggest is facilitated by liquid condensate formation on the plasma membrane by endocytic pioneer proteins, including EPS15 and FCHO^6,13^. While DNM2 is well known for its crucial role in vesicle scission during the final step of CME, our genome-edited DNM2-TagGFP2 faithfully reflects its recruitment to CME sites in two phases, low-level recruitment during the assembly stage followed by a short burst of high-level recruitment during the late stage of CME^20,22,23^. Thus, the AP2M1-TagGFP-T and DNM2-TagGFP2 genome-edited human stem cells used in our earlier studies can elucidate the progression of various CME stages with a temporal resolution of seconds.

Although ITSN1 was proposed to arrive at CME sites with initiation factors EPS15 and FCHOs^6^, knocking it down resulted in an increased rate of CME site initiation and a decreased rate of CME site maturation^18^. To determine the specific stage of CME at which ITSN1 functions, we first investigated the timing of its recruitment. We transiently expressed HaloTag-ITSN1-S in genome-edited cells (AP2-TagRFP-T, DNM2-TagGFP2) through plasmid transfection. Although we observed co-localization of ITSN1 with AP2 and DNM2 on the plasma membrane, indicative of its recruitment to CME sites, CME dynamics in ITSN1-overexpressing cells markedly differed from adjacent control cells without ITSN1 overexpression (Figure s1A). This observation indicates that ITSN1 overexpression perturbs CME and should be avoided when studying its recruitment dynamics.

Therefore, to investigate ITSN1 recruitment timing in a close-to-native context, we generated a genome-edited AP2-TagRFP-T, mEGFP-ITSN1 hiPSC line (Figure 1B and Figure s1B). Endogenously labeled ITSN1 was mainly expressed as the short splicing isoform in hiPSCs (Figure s1B) and, as expected, was recruited without perturbing CME (Figure s1C). We observed that ITSN1 is recruited to CME sites either concurrently with AP2 or shortly thereafter (within 10 seconds); however, it is occasionally recruited significantly later than AP2 (more than 10 seconds). (Figure 1C).

To comprehensively investigate the spatial dynamics of ITSN1 at CME sites, we applied Principal Component Analysis (PCA) clustering to the AP2-positive sites computationally tracked by the cmeAnalysis program^24^ in time-lapse images of genome-edited cells expressing tagged AP2 and ITSN1. The AP2 tracks were classified into five clusters (Figure 1D). Structures in clusters 0-3 have low AP2 or ITSN1 intensity and consist of short-lived events (AP2 lifetimes: 2.2+/-0.6, 6.6+/-5.4, 2.2+/-0.8, 10.9+/-9.9 seconds respectively) that move quickly in X and Y and are unlikely to be authentic CME events^18,25^(Figure 1E, Figure s1D). Cluster 4 comprises high AP2 and ITSN1 intensity structures, which are long-lived (AP2 lifetime: 63.3+/-34.7 sec) and move more slowly in X and Y than the sites in clusters 0-3 (Figure 1E, Figure s1D). Tracks with lifetimes longer than 20 seconds have been considered valid CME events^1,25,26^. Combining the AP2 intensity and lateral movement information, we propose that Cluster 4, enriched for high-ITSN1-intensity structures, represents authentic CME events. Moreover, Cluster 4 tracks with longer AP2 lifetimes recruit higher amounts of ITSN1 and AP2 (Figure 1F), consistent with a high correlation between ITSN1 and AP2 recruitment levels (Figure s1E). In most Cluster 4 tracks, ITSN1 arrives at CME sites shortly after AP2 (Figure 1G): approximately 20% appear before AP2, while more than 60% appear within 10 seconds of the first AP2 detection.

Interestingly, ITSN1 starts disappearing from CME sites before AP2, suggesting that ITSN1 is less likely to be involved in the late stages of CME (Figure 1C, F).

The early-to mid-stage arrival timing of endogenously expressed ITSN1, coupled with the high correlation between ITSN1 and AP2 intensity at CME sites, are more consistent with a role for ITSN1 in CME site stabilization than in site initiation.

### ITSN1 promotes CME site stabilization

To investigate ITSN1 function in CME, we conducted knockdown experiments using transiently expressed shRNAs targeting ITSN1. Each of two different shRNAs resulted in a pronounced reduction of ITSN1 expression (Figure s2A). Our studies used a combination of the two shRNAs for ITSN1 knockdown in genome-edited cells expressing fluorescent protein-tagged AP2 and DNM2. Kymograph analysis revealed a reduction in low-level DNM2 recruitment during the early to middle stages of CME in ITSN1 KD cells, but not in the rapid, pronounced high-level recruitment late in CME (Figure 2A).

**Figure 2.**
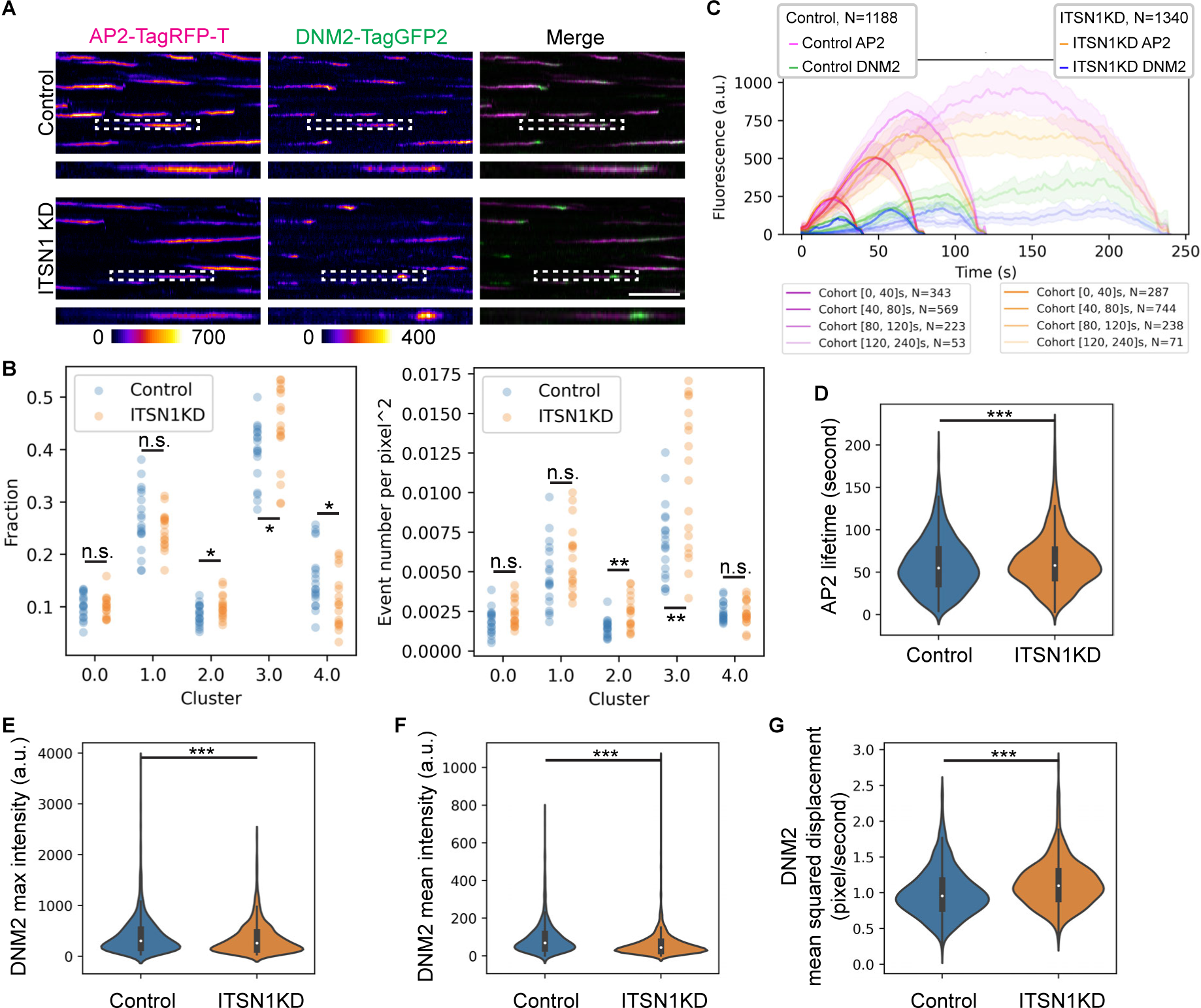
ITSN1 promotes CME site stabilization. (A) Representative kymographs of genome-edited AP2M1-tagRFP-T (magenta) and DNM-tagGFP2 (green) at CME sites in cells 2 days after transfection of plasmids expressing scramble (control) or ITSN1 shRNAs (ITSN1KD). The highlighted kymograph is boxed by a dashed line. Scale bar: 60 sec. (B) Quantification of fraction (left) and event density (right) of each cluster in the valid tracks detected in control and ITSN1KD cells. Each data point represents a dataset collected from one region of interest of TIRF movies. Control: N=19, ITSN1KD: N=18. Statistics: Welch’s t-test, n.s.: not significant, *: p<0.05, **: p<0.01. (C) Averaged intensity vs time plots of cohorts of Cluster 4, DNM2-positive tracks of control and ITSN1KD cells. Tracks are grouped into cohorts by the lifetimes of AP2. The total number of tracks is shown above the plots, and the number of tracks in each cohort is shown below the plots. Error bar: ¼ standard deviation. (D-G) Quantification of AP2 lifetimes and different DNM2 features of Cluster 4 tracks in control and ITSN1KD cells in violin plots. The median is shown as a white dot; the thick vertical line extends from the first quartile to the third quartile of the data, and the thin vertical line extends 1.5x the interquartile range (IQR) from the end of the thick line. Control: N=1180, ITSN1KD: N=1340. Statistics: Welch’s t-test, ***: p<0.005.

To compare CME dynamics in the control and ITSN1KD cells in an unbiased, high-throughput manner, we performed computational analysis of AP2 tracks in AP2M1-TagGFP-T and DNM2-TagGFP2 genome-edited cells. Similar to the analysis in Figure 1D, and as reported previously^22^, AP2 tracks were classified into five clusters, with Cluster 4 representing DNM2-positive CME events (Figure s2B). Comparison of the distribution of AP2 tracks in these clusters in control and ITSN1 knockdown (KD) cells showed a pronounced reduction in the fraction of events grouped in Cluster 4, the authentic CME events, and increased fractions of events classified into Clusters 2 and 3(Figure 2B). Additionally, there was an increase in the total number of events, which did not correlate with a detectable decrease in the density of Cluster 4 events, suggesting that in the ITSN1 KD cells, there may be more small AP2 clusters and/or aborted events (Figure 2B). These data are consistent with the increased initiation density detected in ITSN1KD cells in a previous large-scale study^18^.

Furthermore, when comparing the recruitment dynamics of DNM2 and AP2 at CME sites, we observed reduced DNM2 and AP2 recruitment in the long-lifetime cohorts (>80s). In contrast, the short-lifetime cohort (<80s) showed only moderately reduced recruitment of DNM2 during the early to middle stages, with no noticeable change in the late stage (Figure 2C). These findings suggest that a high level of ITSN1 is particularly crucial for the efficient recruitment of endocytic proteins to long-lifetime CME sites, typically characterized by the recruitment of higher levels of endocytic proteins (Figure 1F and 2C).

Moreover, in ITSN1 KD cells, we observed that AP2 lifetimes were considerably longer (62.6+/-31.2 sec vs 59.0+/-31.15 sec) (Figure 2D), DNM2 intensities at CME sites were lower (Figure 2E, F), and DNM2 lateral movement was increased (Figure 2G). However, AP2 intensity and lateral movement were essentially unchanged (Figure s2C), establishing that DNM2 recruitment to, and stabilization at, CME sites are specific defects.

Together, these results indicate that ITSN1 plays crucial roles in CME site stabilization, particularly through early-to middle-stage DNM2 recruitment.

### ITSN1-S is sufficient to coalesce multiple endocytic proteins into puncta

To define the CME protein interaction network centered around ITSN1, we artificially relocated ITSN1 to the mitochondrial outer membrane by ectopically expressing ITSN1 conjugated to the Tom70 mitochondria targeting peptide.

This relocalization was sufficient to recruit multiple endocytic proteins, including EPS15, FCHO2, AP2M1, DNM2, and EPN1 (Epsin1), to mitochondria and to coalesce them into puncta (Figure 3). These proteins exhibited high molecule exchange rates within the puncta, as demonstrated by Fluorescence Recovery After Photobleaching (FRAP) assay. Both DNM2 and AP2 recover to approximately 60% intensity in less than 1 minute (Figure s3A, B).

**Figure 3.**
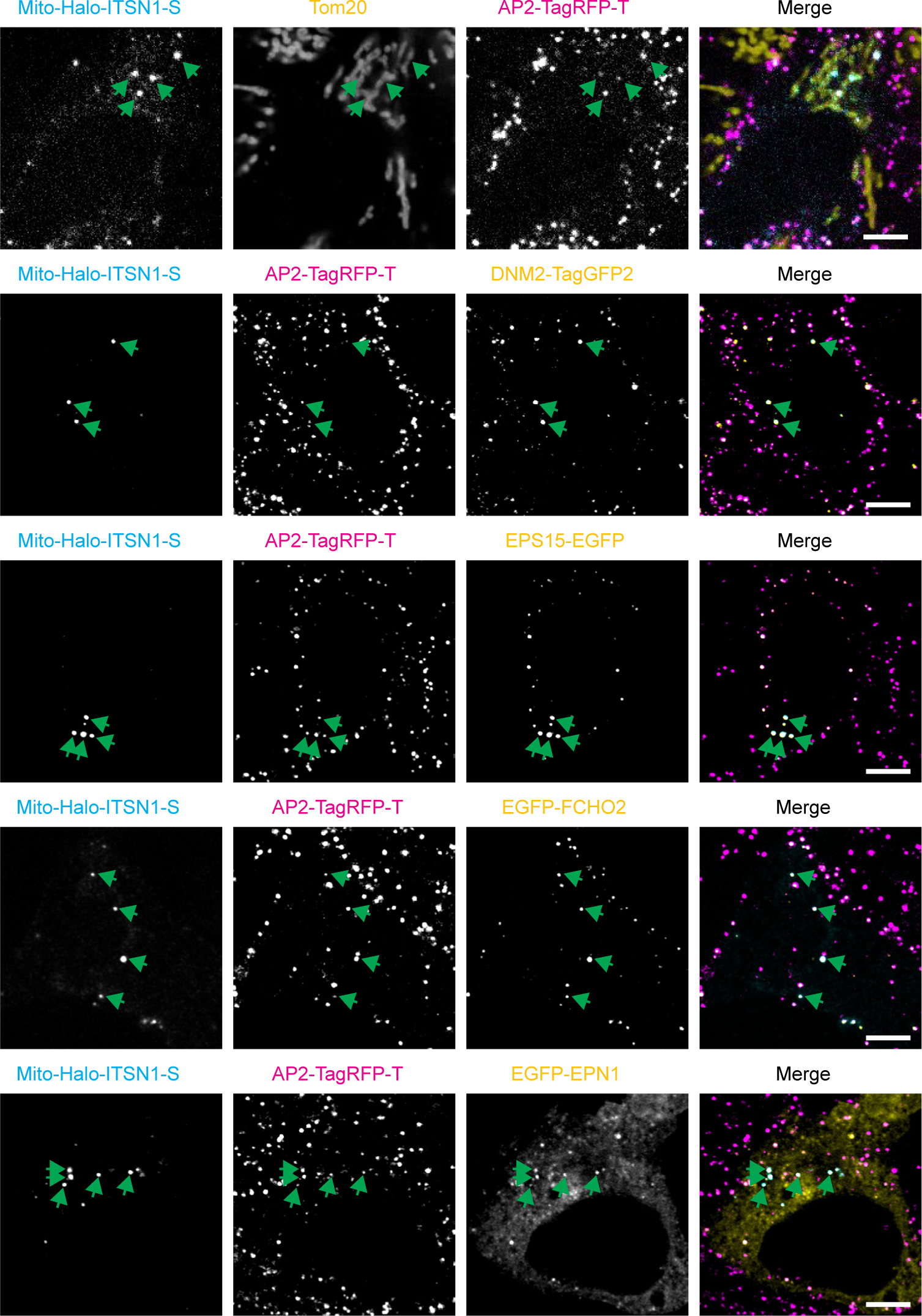
INSN1 recruitment to mitochondria is sufficient to coalesce multiple endocytic proteins into mitochondria-associated puncta. Representative cells fixed 2 days after ectopic mito-HaloTag-ITSN1-S transfection. Images are z-stacks generated by mean intensity projection of 11 optical sections acquired using a Zeiss Airyscan fluorescent microscope. AP2M1 and DNM2 were endogenously tagged with TagRFP-T and TagGFP2, respectively. Tom20 was labeled by immunofluorescence. Plasmids expressing fluorescent protein-conjugated EPS15, FCHO2, or EPN1 were co-transfected with the mito-HaloTag-ITSN1-S expressing plasmid. Scale bar: 5 µm.

EPS15 and its yeast homolog, Ede1, can form condensates when overexpressed^9,27^. To test whether the proteins assembled around relocated ITSN1-S are distinct in composition from those present in EPS15-induced condensates, we artificially targeted EPS15 to mitochondria using the same strategy. Indeed, EPS15 forms puncta that contain FCHO2, AP2, and EPN1.

However, we did not detect DNM2 in the mito-EPS15 puncta (Figure s3C). This result indicates that mito-ITSN1-S assemble puncta containing components beyond those present in the EPS15-FCHO-containing puncta.

Moreover, when we co-expressed mito-ITSN1-S and mito-EPS15 and examined their localization at high resolution, we observed that these proteins localize to the same puncta but occupy distinct regions within the puncta (Figure s3D). This observation further supports the notion that ITSN1 is centered within a protein network distinct in composition from that present in the EPS15-containing initiation condensate.

These data are consistent with the possibility that ITSN1 forms a distinct protein complex and forms a bridge with the EPS15 interaction network, which functions in the initiation of CME sites, and that ITSN1 is responsible for the stable recruitment of proteins such as DNM2 that participate in later stages of CME.

### ITSN1’s different domains play distinct roles in establishing its interaction network

ITSN1 is a multidomain scaffold protein, with its short isoform containing two EH domains, a coiled-coil domain, and five SH3 domains^16^. The EH domains are known to interact with NPF motif-containing proteins, including EPN1^28,29^; the coiled-coil (CC) domain is crucial for ITSN1 oligomerization and its interaction with other CC domain-containing proteins, such as EPS15^29^; and the five SH3 domains interact with proteins containing proline-rich regions, including DNM2^29,30^ (Figure s4A). To dissect the roles of ITSN1’s different domains in establishing its interaction network, we expressed various truncated forms of ITSN1-S (Figure s4A) in the ITSN1KO cells. The ITSN1 KO cells were generated from the AP-TagRFP-T, mEGFP-ITSN1 cells using CRISPR/Cas9 genome-editing and GFP-negative cells were selected by the flow cytometry.

The CC and EH domains are both crucial for ITSN1 recruitment to CME sites (Figure 4A). Mutants lacking either the CC or EH domains could still form puncta when artificially targeted to mitochondria (Figure s4B). However, the CCΔ mutant did not recruit the CME initiation components EPS15 and FCHO2. This observation suggests that ITSN1 is recruited to the initiation site which contains EPS15 and FCHO2 through its CC domain (Figure 4B).

**Figure 4.**
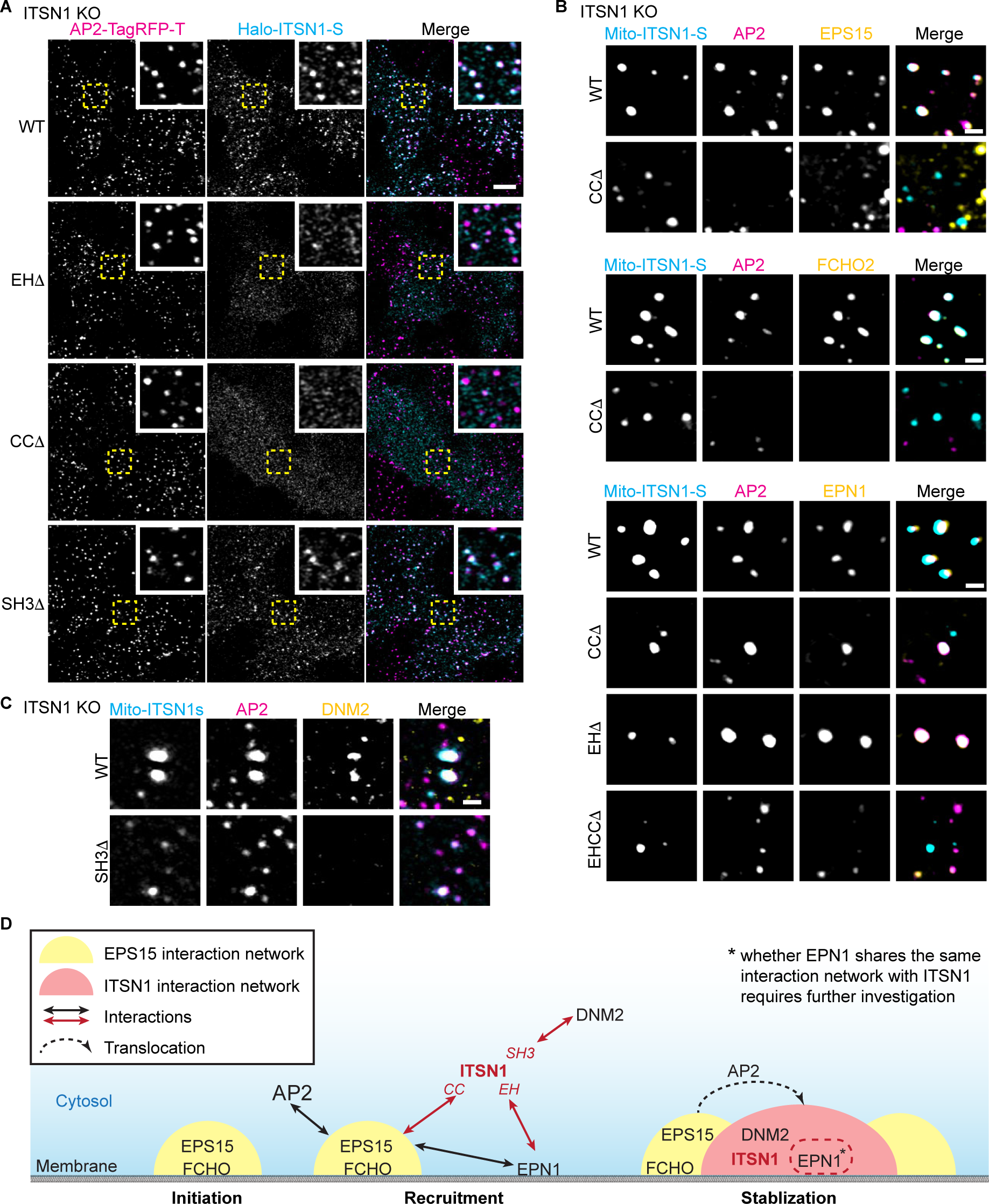
ITSN1’s domains play distinctive roles in establishing its interaction network. (A) Representative cells fixed 2 days after transfection with plasmids expressing wild-type or a truncated form of mito-HaloTag-ITSN1-S. Images were acquired using a Zeiss Airyscan fluorescence microscope. Single-stack images of the basal plasma membrane of cells, where CME sites are labeled by AP2M1-TagRFP-T, are shown. Scale bar: 5 µm. (B, C) Representative images of puncta assembled by mitochondrial recruitment of wild-type or truncation forms of mito-HaloTag-ITSN1-S. Cells were fixed 2 days after plasmid transfection. AP2M1 was endogenously tagged with TagRFP-T. Plasmids expressing fluorescent protein-conjugated EPS15, FCHO2, or EPN1 were co-transfected with the mito-HaloTag-ITSN1-S expressing plasmid (B). DNM2 was labeled by immunofluorescence (C). Scale bar: 1 µm. (D) A schematic model for how ITSN1 is recruited to CME sites, and how it stabilizes CME sites by forming an interaction network among endocytic proteins, including DNM2 and EPN1, which associates with the EPS15-FCHO-containing initiation condensate.

The mitochondria relocalization assay also revealed that CCΔ or EHΔ mutants of ITSN1-S can still interact with EPN1. However, an EHΔCCΔ mutant lacking both the CC and EH domains did not recruit EPN1 to the ITSN1-S-containing puncta (Figure 4B). This result suggests that the interaction between ITSN1-S and EPN1 is mediated redundantly through ITSN1’s CC and EH domains. The most likely explanation for this observation is that ITSN1’s EH domains can interact with EPN1’s NPF motif, while its CC domain mediates an indirect interaction with EPN1 through EPS15 (EPS15 contains EH domains that may interact with EPN1’s NPF motif).

Interestingly, the EHΔCCΔ mutant can still form puncta and recruit DNM2 (Figure s4C). This result supports the conclusion that ITSN1 interacts with DNM2 independently of EPS15/FCHO2 and EPN1.

In contrast, the SH3Δ mutant of ITSN1-S forms dimmer puncta when targeted to mitochondria (Figure s4B). The mito-ITSN1 SH3Δ puncta contain EPS15, FCHO2, EPN1, and AP2M1 (Figure s4D) but do not contain detectable DNM2 (Figure 4C). This result suggests that ITSN1’s SH3 domains are crucial for its interaction with proline-rich domain-containing proteins like DNM2, and that this interaction is important for ITSN1’s ability to establish strong interaction networks.

The above observations are consistent with a model in which ITSN1 is recruited to nascent CME sites through its interaction with a complex of endocytic proteins responsible for initiation. It has been suggested that these early endocytic proteins form a condensate. We propose that subsequent to CME site initiation, ITSN1 mediates a CME site stabilization step by nucleating and stabilizing a protein interaction network associated with, but distinct from, the initiation complex (Figure 4D). Recent data raise the intriguing possibility that both complexes might have properties of liquid condensates.

## Discussion

In this study, we used genome editing to minimize perturbation of the recruitment dynamics of ITSN1, and redefined ITSN1’s function, implicating it in CME site stabilization rather than the previously proposed initiation stage^6^ (Figure 4D). Our results resolve a contradiction between the identification of ITSN1 as a part of the EPS15-FCHO-ITSN1 initiation complex, the observation that ITSN1 has a distinct localization pattern from EPS15 and FCHO at CME sites^12^, and ITSN1 knockdown phenotypes that differ from those of EPS15 or FCHO^18^. Our study highlights the problems associated with overexpressing proteins of interest, especially those with multivalent interaction domains and prone to condensate formation upon overexpression. Overexpressing one or more endocytic proteins can lead to their premature recruitment or delayed disassociation from CME sites, consequently over-stabilizing CME sites^18^.

Here, we showed that ITSN1 interacts with multiple endocytic proteins and contributes to their efficient recruitment to CME sites (Figure 2, Figure 3). We found that low-level, early to middle-stage CME site recruitment of DNM2, prior to its late-stage, high-level recruitment, is particularly dependent on ITSN1 (Figure 2 A, C). Previous studies suggested DNM2 plays roles during multiple stages of CME^31^. While DNM2’s role in scission at the late stage of CME has been extensively studied, its function in the early stage remains understudied and not fully elucidated^26,32^. It was suggested previously that ITSN1 might recruit early-stage DNM2, although no direct evidence was provided^31^. From our findings, we hypothesize that early-stage DNM2, together with other PRD-containing proteins and ITSN1, contributes to CME site stabilization and maturation^26^ by formation of a second liquid condensate after the initiation condensate, through PRD-SH3 multivalent interactions (Figure 4D). We could not gather sufficient evidence that the mito-ITSN1-S-containing puncta are liquid condensates. However, even if we could have demonstrated condensate formation, the uncontrolled expression of mito-ITSN1 prevented us from reconstituting an ITSN1-DNM2 interaction network with the same protein stoichiometry present at natural CME sites. Therefore, despite providing new insights into the previously proposed notion that multiple, distinct liquid condensates form at CME sites^13^, additional evidence is required to determine if and how liquid condensates contribute to assembly of different functional subdomains of CME sites.

ITSN1 has a homolog, ITSN2, which shares the same domain structures with ITSN1. Due to the low expression of ITSN2 in iPSCs, we were not able to genome-edit ITSN2 and compare its dynamics and function to those of ITSN1. However, in the future, it will be important to explore the function of ITSN2 in a cell type in which it is expressed at higher levels.

Our computational analysis showed that in ITSN1 knockdown cells the CME sites with the highest AP2 intensity are absent from the population, suggesting that ITSN1 may be particularly important when a high level of AP2-mediated cargo uptake is required (Figure 2C). Understanding a cargo/signal-specific ITSN1 function may have important implications for its normal function and disease states, as growing evidence suggests that mutations in ITSN1 are linked to increased autism risk, and ITSN1 may play an important role in neuronal development^33–36^.

Although we were able to implicate specific ITSN1 domains in the specific functions (Figure 4), complexity and redundancy in functions and interactions prevented us from pinpointing the domains of ITSN1 that are responsible for its direct interactions with EPN1, FCHO2, and AP2. Additionally, whether EPN1 exists in the same interaction network with ITSN1 or forms a distinct liquid condensate requires further investigation.

In summary, we provided evidence that ITSN1 promotes CME site stabilization. Our results are consistent with the recently proposed model in which CME site formation involves multiple, distinct but interacting protein liquid condensates, and offer insights into how ITSN1 might contribute to the organization of such a multi-condensate system by promoting the stable assembly of the ITSN1-centered condensate and bridging it to an earlier-arriving initiation phase condensate composed of EPS15 and FCHO proteins.

## Acknowledgments

MJ was funded by American Heart Association Postdoctoral Fellowship (18POST34000029). DGD was funded by NIH MIRA grant R35GM118149. pEGFPC1-EPSIN1 was a gift from Pietro De Camilli (Addgene plasmid # 22228 ; RRID:Addgene_22228). The EPS15-EGFP (human) plasmid was a gift from the Taraska Lab at NIH. The EGFP-FCHO2 (mouse) plasmid was a gift from the McMahon Lab at the MRC Laboratory of Molecular Biology. The authors would like to thank Amy Yan and Yuang Song for their assistance in generating plasmids and genome-edited cell lines; the Conklin Lab at UCSF for providing the WTC10 human iPSC line; the Lavis Lab at Janelia Research Campus for providing JF635 HaloTag ligand; the Xu Lab at UC Berkeley for providing the Tom20 antibody; UC Berkeley QB3 MacroLab for purified *S. pyogenes* NLS-Cas9; the Luo Lab at UC Berkeley for sharing their electroporator; the Brar and Ünal Lab at UC Berkeley for sharing their LICOR imager and the UC Berkeley Cancer Research Laboratory Flow Cytometry Facility for iPSC sorting. The authors gratefully acknowledge the developers of cmeAnalysis program and the Python packages and employed in this study for generously providing their code on a free and open-source basis.

## Author contributions

Conceptualization, M.J. and D.G.D.; Methodology, M.J., Y.I. and C.S.; Software, M.J., Y.I. and C.S.; Investigation, M.J. and Y.I.; Resources, M.J., C.S. and D.G.D.; Writing-Original Draft, M.J.; Writing-Review&Editing, M.J., Y.I. and D.G.D.; Visualization, M.J., Y.I. and C.S.; Supervision, D.G.D. and M.J.; Funding Acquisition, M.J. and D.G.D.

## Declaration of interests

The authors declare no competing interests.

## Figure legends

**Figure s1.**
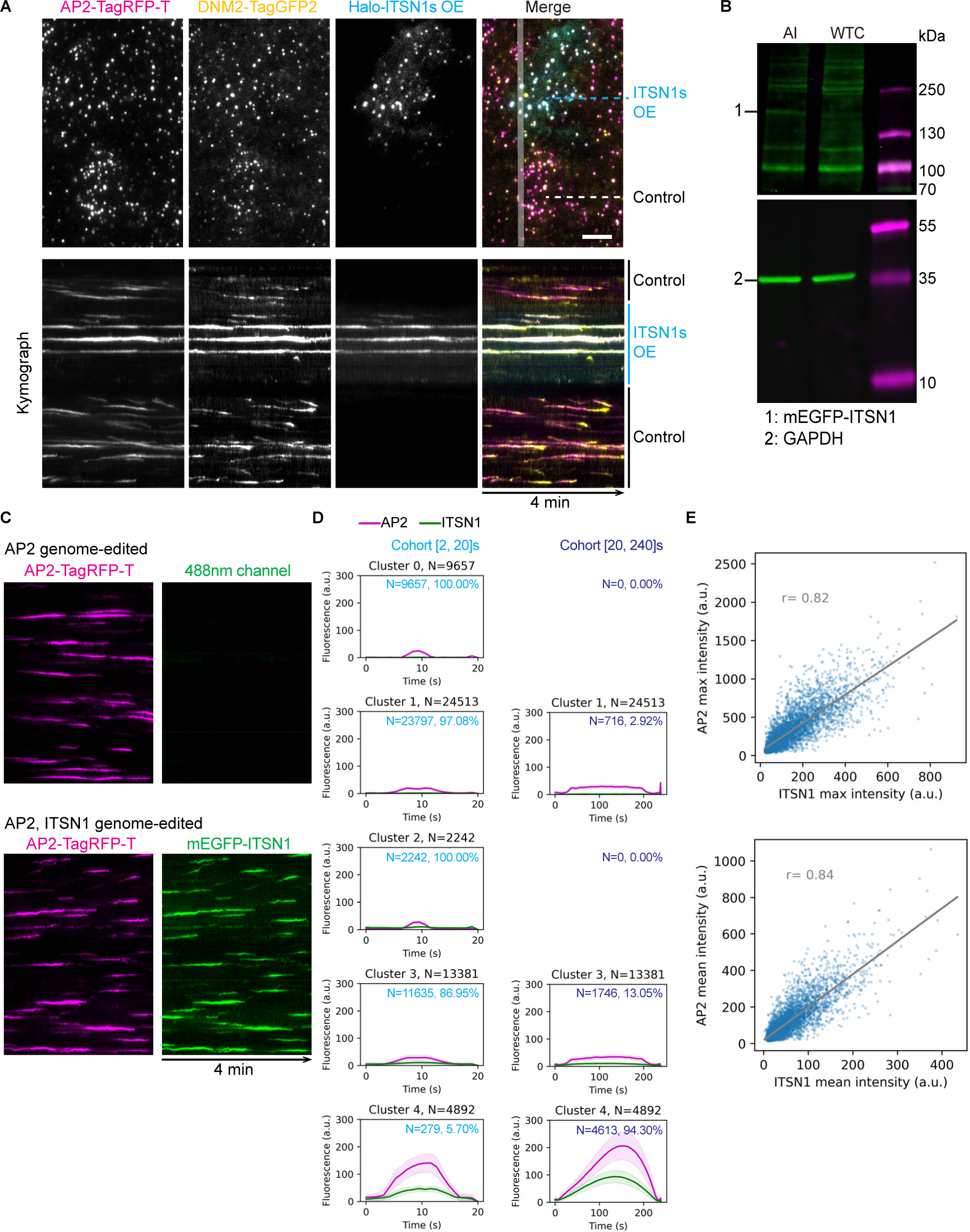
Dynamics of overexpressed or endogenously tagged ITSN1 at CME sites. Related to Figure 1. (A) Top: A representative single-frame image of a TIRF movie of basal plasma membranes of genome-edited, AP2M1-tagRFP-T (magenta), DNM2-tagGFP2 (yellow) human iPSCs with or without significant overexpression of HaloTag-ITSN1 (cyan, labeled by JF635-HaloTag ligand^37^). Scale bar: 5 μm. Bottom: Kymographs of CME sites highlighted in the top panel. (B) Immunoblot analysis of cell extracts from genome-edited (AP2M1-tagRFP-T, mEGFP-ITSN1; AI) and control (WTC) human iPSCs. The labeled proteins were detected with Tag(CGY)FP and GAPDH (loading control) antisera, respectively. (C) Representative kymographs of TIRF movies of basal plasma membranes of genome-edited AP2M1-tagRFP-T (magenta) and AP2M1-tagRFP3-T (magenta), mEGFP-ITSN1 (green) human iPSCs. (D) Cohort (0-20 seconds and 20-240 seconds) plots of AP2 events from each cluster. Cluster 4 represents ITSN1-positive events where a strong ITSN1 signal is detected. Data are presented as mean values +/- 1/4 standard deviation. The total number of tracks in each cluster is shown on top of the plot, and the number of tracks in each cohort is shown next to the plot. Please note that Cluster 0 and 2 don’t have any tracks in the 20-240 cohort. (E) Scatter plots of maximum (top) and average (bottom) intensities of ITSN1 and AP2 of Cluster 4 tracks. Each data point represents a track. The linear regression line is shown in grey. R: correlation coefficient. N=4892.

**Figure s2.**
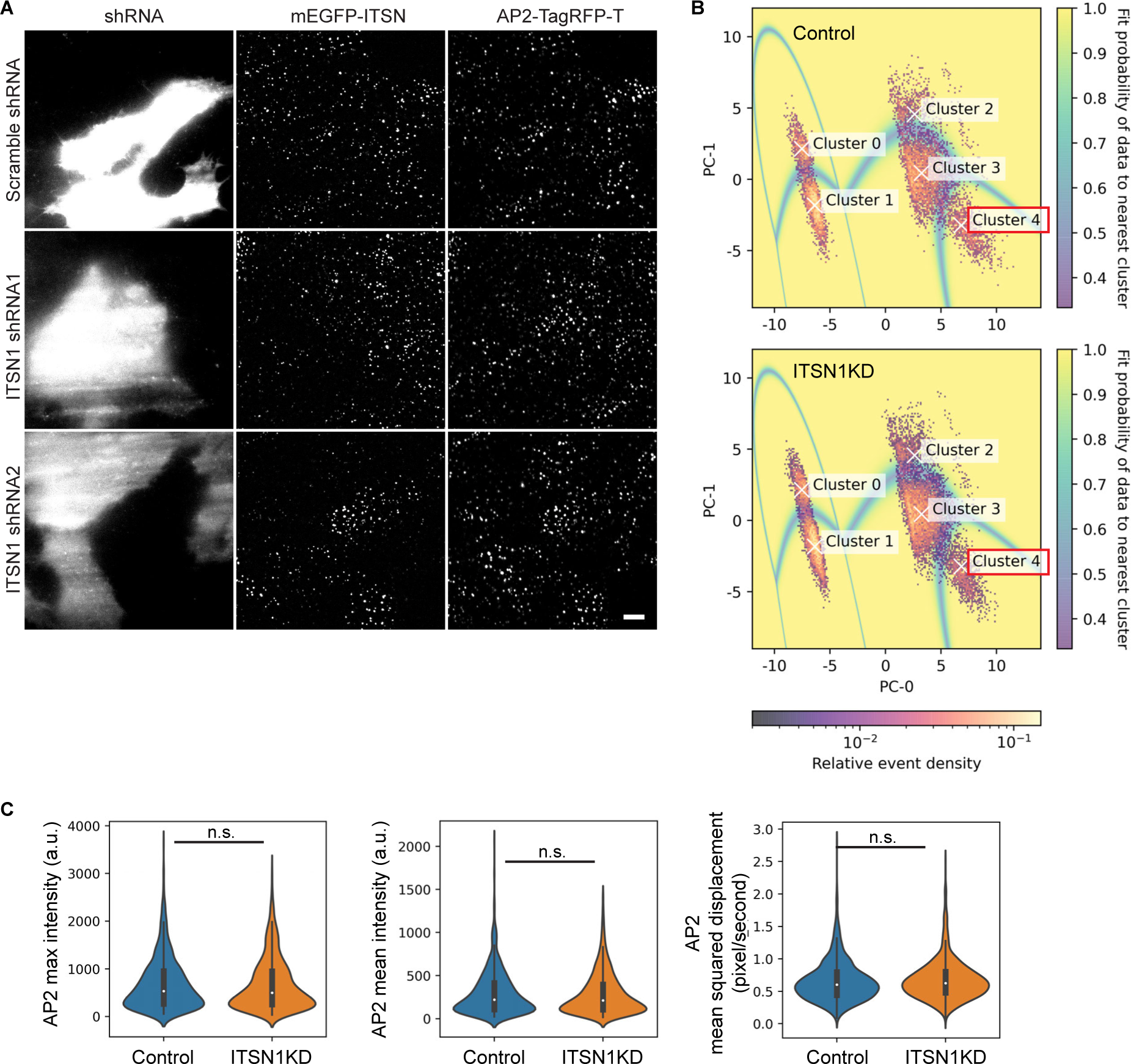
ITSN1 promotes CME site stabilization, related to Figure 2. (A) Representative TIRF micrographs of genome-edited AP2M1-tagRFP-T, mEGFP-ITSN1 iPSCs two days after transfection with plasmids co-expressing TagBFP and scramble or ITSN1 shRNAs. Scale bar: 5 µm. (B) 2-D histogram of the first two principal components (PCs) of AP2 and DNM2 dynamic features in control (top) and ITSN1KD (bottom) cells. Valid tracks detected by CMEanalysis^24^ were used to generate filtering methods and for subsequent analysis. Relative event density shows the distribution of the valid tracks in the principal component space and the shaded underlay (fit probability of data to nearest cluster) represents the probability of the simulated data points belonging to the nearest cluster. Cluster 4 shows data points in the DNM2-positive cluster, as previously published. (C) Quantification of different features of Cluster 4 tracks in control and ITSN1KD cells in violin plots. The median is shown as a white dot, the thick vertical line extends from the first quartile to the third quartile of the data, and the thin vertical line extends 1.5x the inter-quartile range (IQR) from the end of the thick line. Control: N=1180, ITSN1KD: N=1340. Statistics: Welch’s t-test, n.s.: not significant.

**Figure s3.**
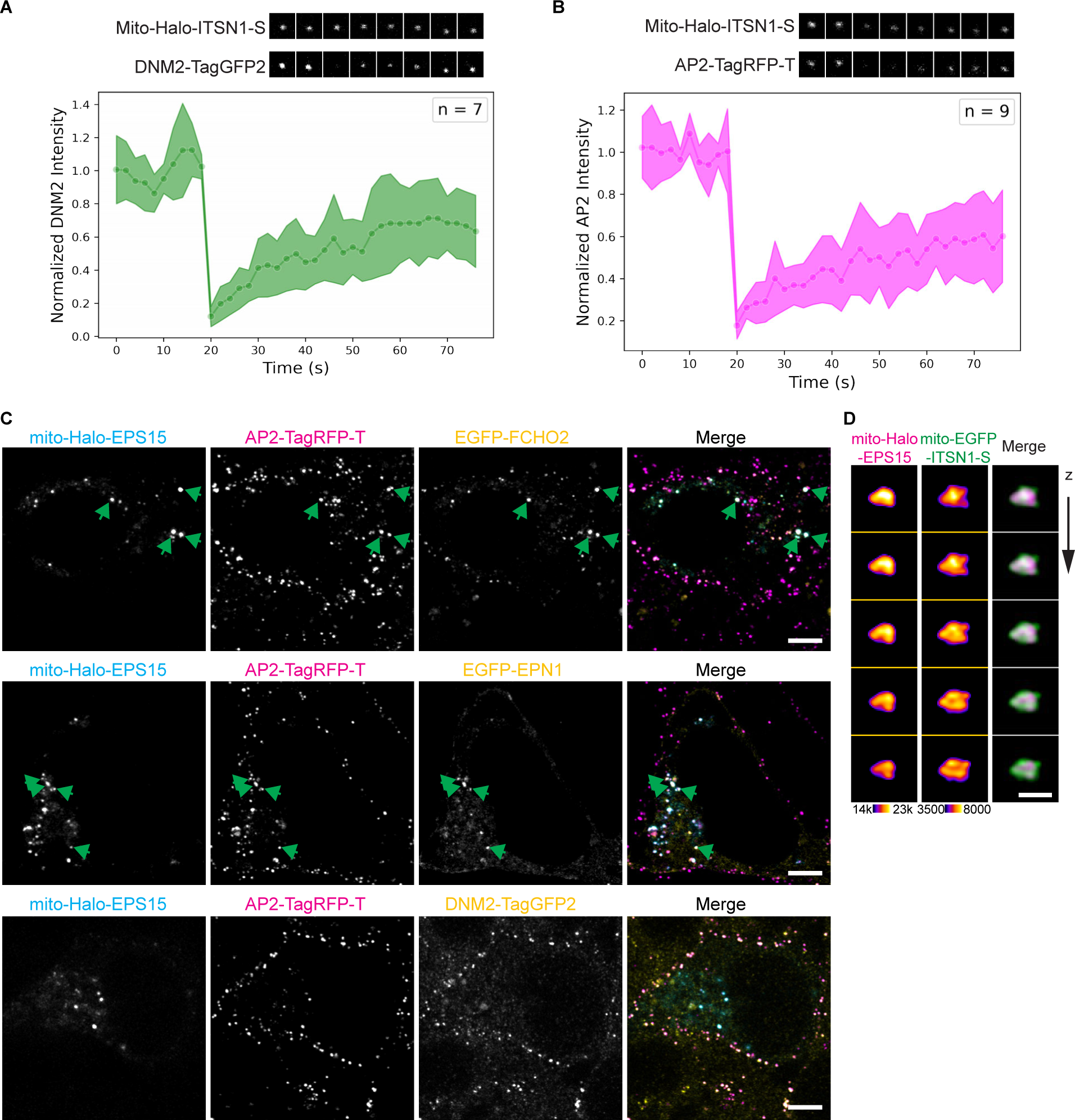
ITSN1 recruitment to mitochondria is sufficient to form puncta with dynamic DNM2 recruitment. Related to Figure 3. (A, B) FRAP assay of mito-ITSN1-S-containing puncta showed rapid recovery of DNM2 (A) and AP2 (B) after photobleaching. (C) Representative cells fixed 2 days after ectopic mito-HaloTag-EPS15 transfection. Images were generated by mean intensity projection of 11 z-stacks acquired by a Zeiss Airyscan fluorescent microscope. AP2M1 and DNM2 were endogenously tagged with TagRFP-T and TagGFP2, respectively. Plasmids expressing fluorescent protein-conjugated FCHO2 or EPN1 were co-transfected with the mito-HaloTag-EPS15 expressing plasmid. Scale bar: 5 µm. (D) Representative large puncta in mito-Halotag-EPS15 and mito-EGFP-ITSN1-S co-expressing cells. Cells were fixed 2 days after transfection of plasmids. Images were generated by Zeiss Airyscan fluorescent microscope. Z-step: 150 nm. Scale bar: 1 µm.

**Figure s4.**
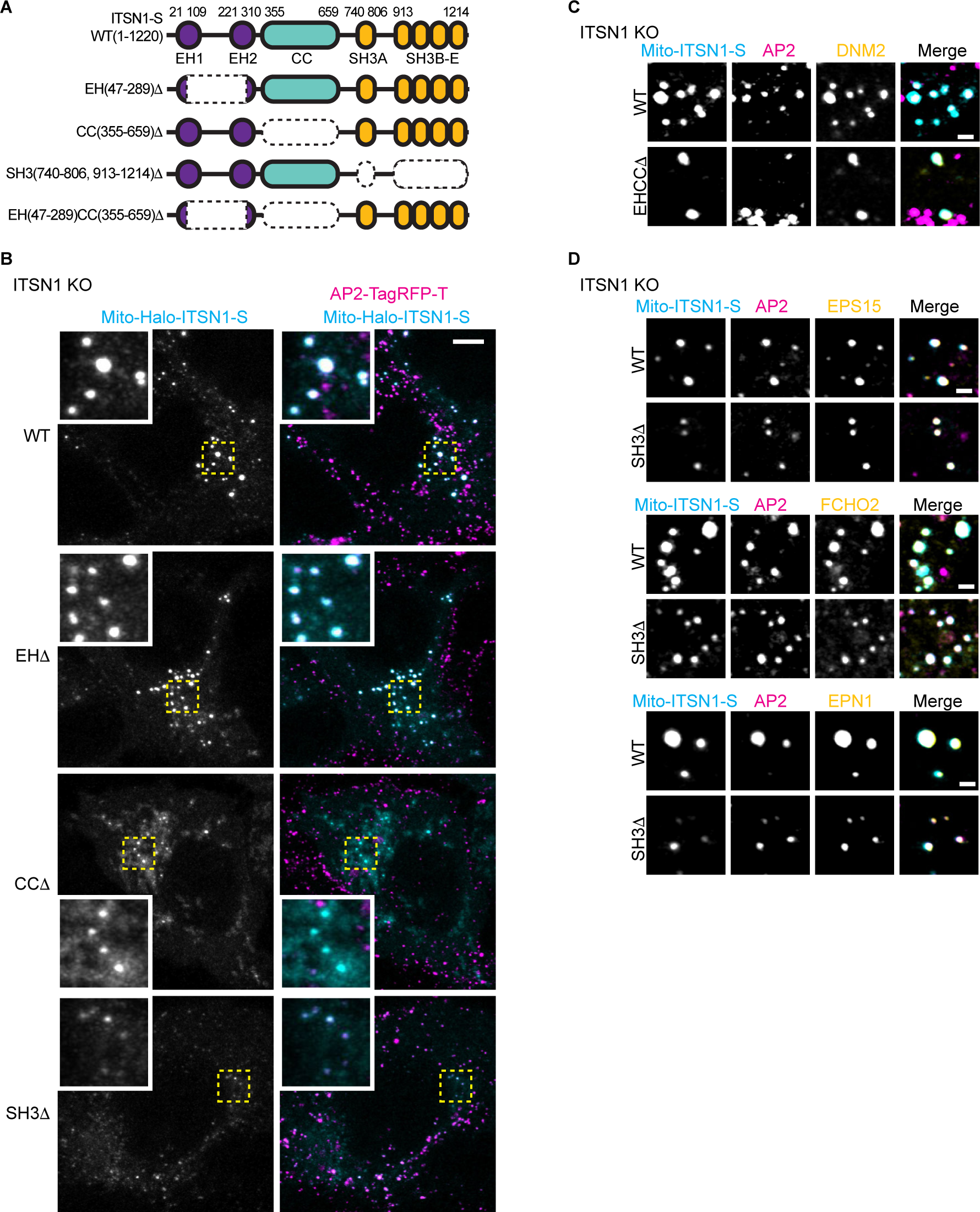
ITSN1’s different domains play distinctive roles in establishing its interaction network. Related to Figure 4. (A) An illustration of domains of wild-type ITSN1-S and the truncation mutants investigated in this study. (B) Representative cells expressing wild-type and truncation forms of mito-HaloTag-ITSN1-S. AP2M1-TagRFP-T, ITSN1KO genome-edited cells were fixed 2 days after plasmid transfection and imaged using a Zeiss Airyscan fluorescence microscope. Scale bar: 5 µm. (C) Representative puncta formed by wild-type or EHCCΔ mito-HaloTag-ITSN1-S. AP2M1-TagRFP-T, ITSN1KO cells were fixed 2 days after plasmid transfection and DNM2 was localized by immunofluorescence. Cells were imaged using a Zeiss Airyscan fluorescence microscope. Scale bar: 1 µm. (D) Representative puncta formed by wild-type or SH3Δ mito-HaloTag-ITSN1-S. Plasmids expressing GFP-conjugated EPS15, FCHO2, or EPN1 were co-transfected to visualize the localization of these endocytic proteins. AP2M1-TagRFP-T, ITSN1KO genome-edited cells were fixed 2 days after plasmid transfection and imaged using a Zeiss Airyscan fluorescence microscope. Scale bar: 1 µm.

## Materials and Methods

### RESOURCE AVAILABILITY

#### Lead contact

Further information and requests for data and materials should directed to and will be fulfilled by the lead contact, David G Drubin.

#### Materials availability

Plasmids and cell lines generated in this study will be available upon request.

#### Data and code availability

Raw data and original code have been deposited at Zenodo. DOIs are listed in the key resources table. They are publicly available as of the date of publication.

### EXPERIMENTAL MODEL AND STUDY PARTICIPANT DETAILS

The WTC hiPSC parental cell line (male, Coriell Cat. # GM25256) was obtained from the Bruce Conklin Lab at UCSF. All the cell lines used in the study were generated by genome editing of the WTC hiPSC line by the authors. AP2, DNM2, and AP2, ITSN1 dual-edited cell lines used in this study were verified by genomic DNA purification, Sanger sequencing and western blotting. All the cell lines used in this study tested negative for mycoplasma contamination by PCR and authenticated by the University of California, Berkeley Cell Culture Facility using short tandem repeat (STR) analysis. The hiPSCs were cultured on Matrigel (hESC-Qualified Matrix, Corning) in StemFlex medium (Thermo Fisher) with Penicillin/ Streptomycin in 37°C, 5% CO2. Cultures were passaged with Gentle Cell Dissociation reagent (StemCell Technologies, Cat#: 100-0485) twice every week.

### METHOD DETAILS

#### Cell Culture

The WTC hiPSC line was sourced from the Bruce Conklin Lab at the University of California, San Francisco. The hiPSCs were grown on Corning Matrigel hESC-Qualified Matrix (Sigma-Aldrich, Cat# CLS354277) in StemFlex medium (Thermo Fisher Scientific, Cat# A3349401) supplemented with Penicillin/Streptomycin at 37°C with 5% CO2. Passaging of cell cultures was conducted biweekly using Gentle Cell Dissociation Reagent (StemCell Technologies, Cat# 100-0485).

#### Genome-editing

The Cas9-sgRNA complex electroporation method was used to knock in mEGFP at the 5’ of the ITSN1 gene in AP2M1-tagRFP-T genome-edited hiPSCs. *S. pyogenes* NLS-Cas9 was purchased from the University of California Berkeley QB3 MacroLab. sgRNA targets AGGTGTTGGAAACTGAGCCA was purchased from Synthego. NEBuilder HiFi DNA Assembly Master Mix (New England Biolabs, Cat# E2621S) was used to construct the donor plasmid containing 450 bp ITSN1 5’ homology arm including start codon, mEGFP, a linker sequence: AAG TCC GGA GGT ACT CAG ATC TCG AGG, and 447bp ITSN1 3’ homology arm sequence. Following electroporation (Lonza, Cat#: VPH-5012) of the Cas9-sgRNA complex and the donor plasmid, mEGFP-positive hiPSCs were individually sorted into Matrigel-coated 96-well plates using a BD Bioscience Influx sorter (BD Bioscience) three days later. The confirmation of clones was carried out through PCR and Sanger sequencing of the genomic DNA surrounding the insertion site. Both alleles of ITSN1 were tagged with mEGFP.

The Cas9-sgRNA complex electroporation method was used to knockout ITSN1 in the AP2M1-tagRFP-T, mEGFP-ITSN1 dual-edited hiPSCs. sgRNA targets GAAGTTCGAGGGCGACACCC was purchased from Synthego. mEGFP-negative and tagRFP-T-positive hiPSCs were pooled into Matrigel-coated plates using a BD Bioscience Influx sorter (BD Bioscience), three days after electroporation (Lonza, Cat#: VPH-5012) of the Cas9-sgRNA complex. Verification of the absence of mEGFP-ITSN1 expression was conducted through fluorescence microscopy.

#### Lysate preparation and Western Blotting

Each iPSC lysate was prepared from one well of a 6-well plate. Cells were washed in PBS, incubated in 200ul lysis buffer (50mM Hepes pH 7.4, 150mM NaCl, 1mM MgCl2, 1% NP40, phosSTOP (Roche, Cat# 4906845001), protease inhibitor (Roche, Cat# 1183617001)) for 15 minutes at room temperature. After collecting the supernatant, it was centrifuged at 4°C and 13,000rpm for 15 minutes. The lysate was combined with Laemmli reducing buffer (5% β-mercaptoethanol) and boiled for 10 minutes for Western blot analysis.

PageRuler Plus Prestained Protein Ladder (Thermo Fisher Scientific, Cat# 26619) and cell lysate samples were loaded onto a polyacrylamide gel for SDS-PAGE and transferred onto nitrocellulose membrane via overnight wet transfer. Blots were blocked with 5% milk in TBS (Tris-buffered saline) and incubated with primary antibody at 4°C overnight. For primary antibody blotting, anti-(CGY)FP (1:2000 dilution in 1% milk in TBS, Evrogen, Cat# AB121) and anti-GAPDH (1:10000 in TBST, Abcam ab9485) were used. Finally, blots were incubated with IRDye 800CW secondary antibodies (Licor, Cat# 926-32211 and Cat# 926-32210) diluted 1:10,000 in TBST containing 5% milk for 1 hour at room temperature and imaged on the LICOR Odyssey DLx imager.

#### TIRF live-cell imaging

Two days prior to imaging, human induced pluripotent stem cells (hiPSCs) were plated onto Matrigel-coated 8-well chambered cover glasses (Cellvis, Cat# C8-1.5H-N). If plasmid transfection was required (ITSN1 overexpression or knockdown), during cell plating, 300ng of plasmid was transfected using Lipofectamine Stem Transfection Reagent (Thermo Fisher Scientific, Cat# STEM00001) in 150µl Opti MEM, along with 10µM ROCK inhibitor Y-27632 (Chemdea, Cat# CD0141) for each well. Four hours post-transfection, 150µl StemFlex medium with 10µM ROCK inhibitor was added to each well, followed by a switch to StemFlex medium 24 hours after transfection. For HaloTag-ITSN1 imaging, before imaging, cells were incubated in StemFlex medium with 100 mM JF635-HaloTag ligand for 1 hour. Unbound ligands were removed through three washes, each involving a 5-minute incubation in prewarmed StemFlex medium. Imaging was conducted using a Nikon Ti-2 inverted microscope equipped with 60x total internal reflection fluorescence (TIRF) optics and a sCMOS camera (Hamamatsu). Cells were maintained at 37 °C throughout the imaging process using a stagetop incubator (OKO Lab) in StemFlex medium supplemented with 10 mM HEPES. Nikon Elements software was utilized for image acquisition, capturing channels sequentially at a 1-second intervals with a 300ms exposure time over a 4-minute duration.

#### TIRF image processing

Three generalized processing steps were applied to cluster AP2-positive events: track feature abstraction, feature dimensionality reduction, and event clustering.

First, trajectories, delineated by fitted positions and intensities for individual AP2 positive events, were created using cmeAnalysis^24^. AP2-tagRFP-T served as the master channel for clathrin-mediated endocytosis in cmeAnalysis tracking experiments. DNM2-tagGFP2 or mEGFP-ITSN1 was used as a secondary channel.

Subsequently, within Python Jupyter notebooks, AP2 and ITSN1(or DNM2) tracks underwent decomposition into dynamic features that characterized the events’ positional changes and brightness variations. Each tracked event, once an arbitrary array of intensities and positions, was now a discrete vector of fixed length. Each track was mapped to discrete features to generalize the dynamics of heterogeneous tracked events into a set of interpretable coordinates.

Following feature abstraction, the output array is a 2-dimensional matrix with N rows (N tracked events) and M columns (M discrete features per track). These features were individually scaled to normal distributions to remove the variability in scale and dampen the effects of outliers. For instance, the ‘lifetime’ feature (AP2 lifetime) ranged from a few seconds to several minutes on a scale of seconds, whereas the ‘DNM2-peak fraction’ feature (where the DNM2 peak is located within one AP2 event) ranges from 0 to 1. Following feature re-scaling, these events, which each contain over thirty features, were projected to a lower-dimensional space via principal component analysis. The derived clusters were separated using a Gaussian Mixture Model and events were assigned to clusters based on their highest probability of identity to one cluster.

The resultant data underwent downstream analysis.

Comprehensive Jupyter notebook codes detailing each step are accessible at http://doi.org/10.5281/zenodo.10988767 as of the date of publication.

#### Confocal fluorescence imaging

Two days prior to fixation, hiPSCs were single-cell passaged onto Matrigel-coated 8-well chambered cover glasses (Cellvis, Cat# C8-1.5H-N). During cell plating, 300ng of plasmid was transfected using Lipofectamine Stem Transfection Reagent in 150µl Opti MEM, along with 10µM ROCK inhibitor Y-27632 (Chemdea, Cat# CD0141) for each well. Four hours post-transfection, 150µl StemFlex medium with 10µM ROCK inhibitor was added to each well, followed by a switch to StemFlex medium 24 hours after transfection.

Before fixation, cells were incubated in StemFlex medium with 100 mM JF635-HaloTag ligand for 1 hour, and unbound ligands were removed through three washes, each involving a 5-minute incubation in prewarmed StemFlex medium. Subsequently, cells were fixed for 20 minutes in 4% (v/v) PFA (Electron Microscopy Sciences, Cat#: 15710) in PBS, followed by three 10-minute washes in PBS.

For samples requiring antibody staining, a 20-minute block was performed using blocking buffer [3% (w/v) BSA and 0.1% (w/v) Saponin in PBS]. Primary antibody immunostaining was performed overnight at 4°C. The next day, samples underwent three 10-minute washes in washing buffer (0.1x blocking buffer in PBS). Secondary antibody incubation took place for 30 minutes at room temperature, followed by three 10-minute washes in washing buffer and three 10-minute washes in PBS.

Fixed cell samples were imaged on a Zeiss LSM 900 inverted microscope equipped with a Zeiss 60x objective (NA1.4) and an Airyscan 2 detector using Multiplex 4Y line scanning mode. Image acquisition and processing were performed using the ZEN 3.1 system. Channels were acquired sequentially, and images underwent 3D Airyscan processing.

#### FRAP assay

Movies for FRAP analysis were acquired using a Zeiss 60x objective (NA1.4) on a Zeiss LSM900 microscope in the point-scan confocal imaging mode. For acquisition, the pinhole size and laser speed were set to max and the optimal frame size was selected. Images were acquired every two seconds and photobleaching was conducted after a 10 frame pre-bleach period. To ensure that the photobleached puncta stayed in focus, we imaged a mito-HaloTag-ITSN1 (labeled by JF635-HaloTag ligand^37^) reference protein that localizes to the puncta but does not photobleach. All imaging was conducted at 37°C.

For FRAP analysis, the acquired movies were first background subtracted and subjected to photobleach correction using the imageJ photobleach correction function (exponential fit). For all puncta analyzed, the reference puncta of the acquired movies were tracked using the imageJ plug-in suite PatchTrackingTools^38,39^, and the intensity change of the bleached channel at the reference puncta location was monitored. The obtained intensity values were pooled by first normalizing every intensity value of a trace to the first measured intensity value, and then to then to the average intensity value over the prebleach period. After normalization, these traces were pooled, and the average trace was plotted along with the standard deviation for each time point. Importantly, in each case, we bleached the entire puncta, so whether the recovery is diffusion or reaction-limited is unclear. Plots were generated in a python notebook using the seaborn, numpy, and pandas packages. The notebook is accessible at http://doi.org/10.5281/zenodo.10988767 as of the date of publication.

### QUANTIFICATION AND STATISTICAL ANALYSIS

Excel, MATLAB and Python were used for quantification and statistical analysis. The statistical details of experiments can be found in each figure and its corresponding legend, where applicable.

### KEY RESOURCES TABLE

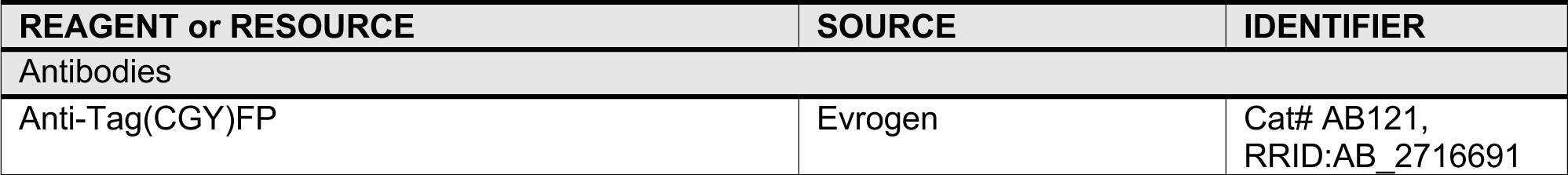
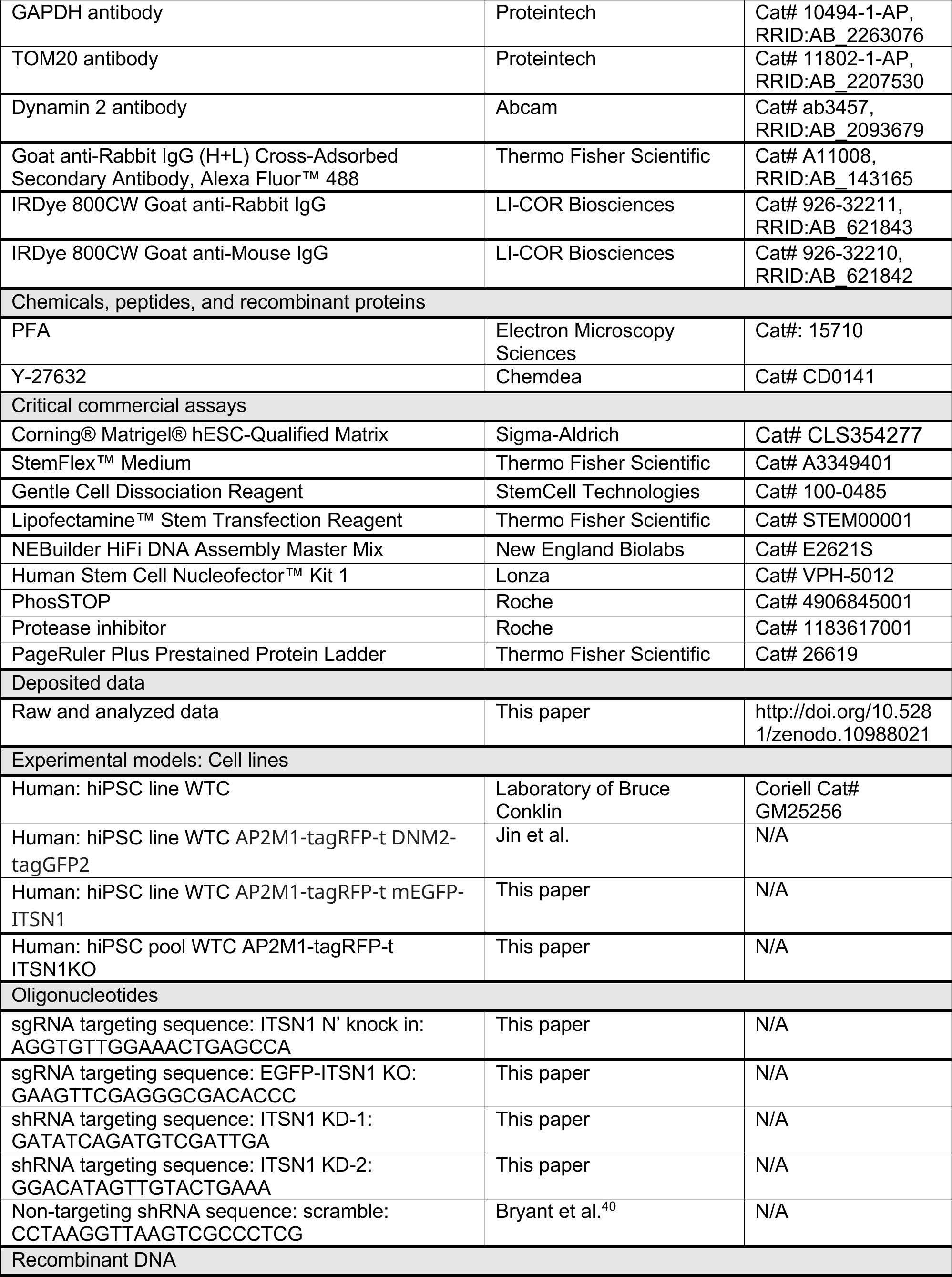
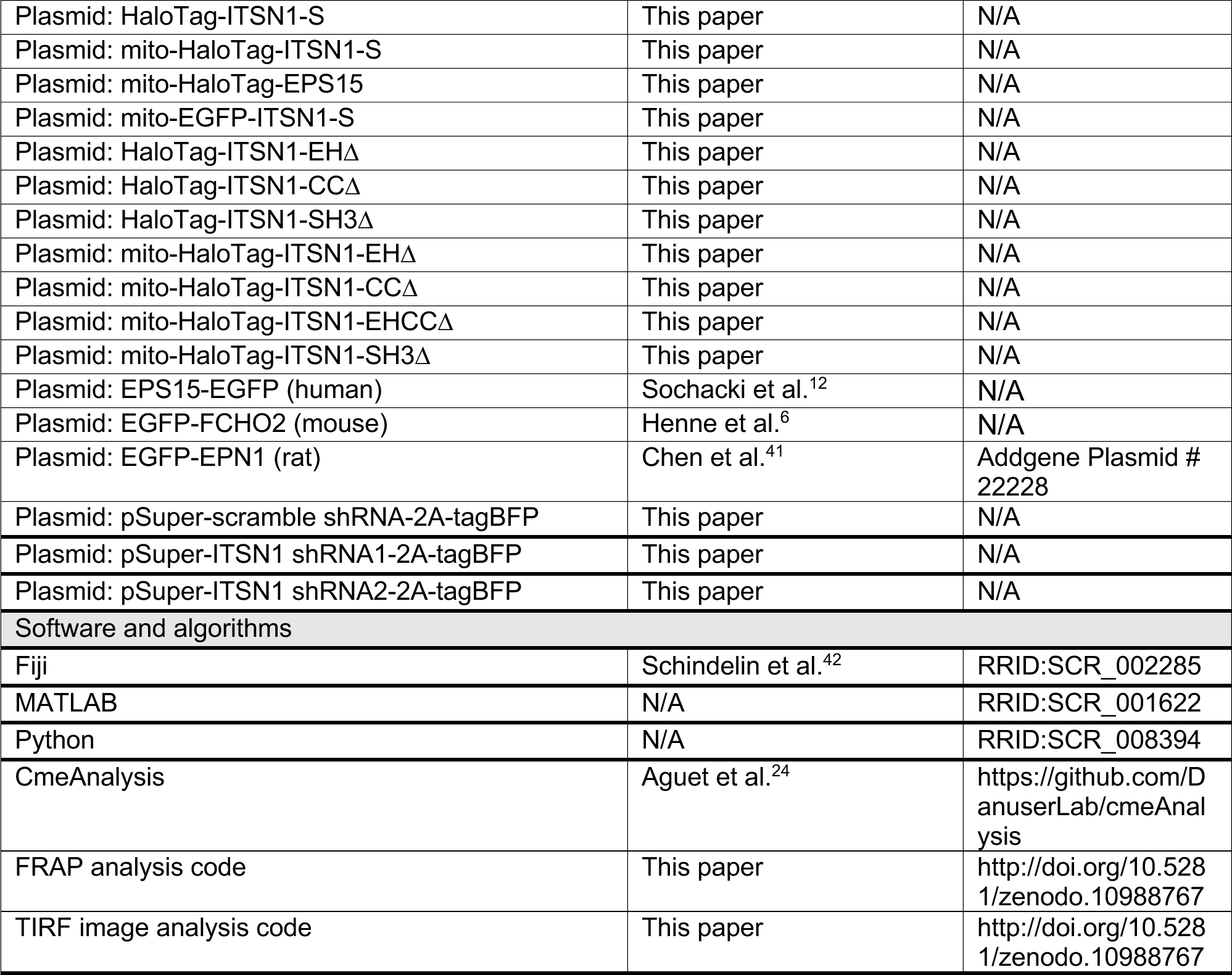

## Notes

http://doi.org/10.5281/zenodo.10988021

http://doi.org/10.5281/zenodo.10988767

### Competing Interest Statement

The authors have declared no competing interest.

